# Discovery of a novel bandavirus using metagenomic sequencing in a retrospective analysis of an unresolved 2020 mortality event involving black vultures in the northeastern United States

**DOI:** 10.1101/2024.12.21.629921

**Authors:** Lusajo Mwakibete, Axel O. G. Hoarau, Vida Ahyong, Eric Waltari, Susan J. Bender, Sherrill Davison, Kevin D. Niedringhaus, Michelle L. Gibison, Roderick B. Gagne, Erica A. Miller, Lisa A. Murphy, Amy Kistler, Cristina M. Tato

**Affiliations:** Chan Zuckerberg Biohub San Francisco, California; Wildlife Futures Program, University of Pennsylvania School of Veterinary Medicine, Kennett Square, PA; Pennsylvania Animal Diagnostics Laboratory System (PADLS), University of Pennsylvania School of Veterinary Medicine, Kennett Square, PA

## Abstract

Investigations of wildlife diseases and mortality events can sometimes lead to inconclusive results due to limitations in diagnostics combined with an ever-increasing number of emerging viruses. The use of tools such as unbiased metagenomic next generation sequencing (mNGS) can facilitate the identification of causative agents where conventional investigation methods fail. We performed a retrospective mNGS analysis on RNA isolated from postmortem samples collected during a black vulture (*Coragyps atratus*, family: Cathartidae) mortality event that occurred in eastern Pennsylvania and western New Jersey in 2020. We describe the discovery and identification of a novel species of bandavirus (*Phenuiviridae* family) in case specimens from this die-off, as well as some of the associated pathological findings. The *Bandavirus* genus comprises tickborne viral species that have been reported across five continents and implicated in outbreaks in a variety of mammalian hosts, including humans, and in avian species making them important potential sources of zoonotic spillover events. Genomic and phylogenetic analysis of the bandavirus detected in this study indicate its closest relative corresponds to Hunter Island virus, a bandavirus previously implicated in albatross mortality events off the coast of Tasmania, Australia. Follow-up PCR testing of samples from additional vultures from the same cohort confirmed that this new bandavirus is the likely cause of death.

## Introduction

Metagenomic next generation sequencing (mNGS) has proven to be an effective tool for the investigation of unresolved wildlife disease events ^1–3^. Employing unbiased approaches, like mNGS, can fill a widening gap in the ability to effectively detect unknown pathogens where targeted approaches fail since no prior knowledge is needed to direct the investigation. This need is underscored by both the limited availability of wildlife diagnostics and the increasing emergence of new zoonotic viruses with epidemic potential over the last 25 years ^4,5^.

Additionally, known viruses, including mosquito-borne pathogens such as Zika virus and Chikungunya virus, as well as tick-borne encephalitis viruses (subtypes European, Far eastern and Siberian), are steadily establishing themselves as endemic across a shifting geographical range due to both environmental and demographic factors, such as urbanization, habitat encroachment, climate change, and population displacement ^6–9^. Thus, as vector-transmitted viruses, members of the Order *Hareavirales* (formerly, *Bunyavirales*) are of particular concern both because of their large host ranges and because they are important causative agents of animal and human hemorrhagic disease outbreaks ^10^.

Some of the more widely known *Hareavirales* include Rift Valley fever virus and Crimean Congo hemorrhagic fever virus. Lesser known species of this order belong to the *Phenuiviridae* family and the tickborne, *Bandavirus* genus, comprised of nine classified species and three related, but unclassified species ^11^. Three species, Bhanja virus (*Bandavirus bhanjanagarense)*, Heartland virus (*Bandavirus heartlandense*) and Dabie bandavirus (*Bandavirus dabieense,* also known as severe fever and thrombocytopenia syndrome virus (SFTSV)) have been described as causing disease in humans ^10,12–16^. Additionally, there is serological evidence implicating a fourth bandavirus species, Guertu virus (*Bandavirus guertuense),* is capable of infecting humans ^17^.

Bandaviruses are enveloped, negative-stranded RNA viruses with a genome consisting of three segments, the L, M and S segments. The L segment encodes the RNA-dependent RNA-polymerase (RdRp), the M segment encodes the glycoprotein precursor, and the S segment contains the nucleocapsid ^10^. In addition, a common characteristic of the S segment is additional ambisense coding of a non-structural protein that is thought to be an important virulence factor ^10^. The segmented genome and wide host range for infection makes bandaviruses particularly suited to genomic reassortment and potential zoonotic spillover.

In this study, we undertook a retrospective analysis of post-mortem samples collected from black vultures (*Coragyps atratus*, family: Cathartidae) to elucidate the cause of an unresolved mortality event that occurred in eastern Pennsylvania and western New Jersey in 2020. Using RNA mNGS we were able to identify a novel species of bandavirus (*Phenuiviridae* family) that was present in tissues of three representative cases. Follow-up PCR testing on additional vultures with similar pathological findings from the same cohort indicates that this new bandavirus is the likely cause of death. Here, we focus on characterizing and contextualizing this new virus within the *Bandavirus* genus and the *Phenuviridae* family.

## Methods

### Description of the Mortality Event

In the summer of 2020 an isolated mortality event occurred affecting black vultures from five counties across eastern Pennsylvania and western New Jersey. In New Jersey, over 20 free-roaming, wild black vultures were found dead on the grounds of a wildlife rehabilitation center (Mercer County) over the course of three weeks in late June to early July, and additional birds were found dead at a fishery (Warren County) in September. Free-ranging wild black vultures were also found dead on the grounds of a zoological facility in central Pennsylvania (Dauphin County), over several weeks in late July, and several were found dead at two other sites in Pennsylvania (Delaware and Dauphin Counties), in September and October. A total of 32 suspected cases were collected from these sites for necropsy and pathological analysis. Out of the 32 birds, two were too decomposed to proceed to necropsy and three were sent to another diagnostic lab. The remaining 27 black vultures presented for necropsy at New Bolton Center, University of Pennsylvania.

### Postmortem Examination

All birds were deceased upon arrival and underwent full gross examination for pathological findings. Sterile dissection scissors were used to remove approximately 5mm^3^ samples of tissue from various organs. Tissue samples were put into cryovials containing DNA/RNA Shield (Zymo Research, CA, USA) before freezing at -18°C. Additional tissues were put directly into formalin at room temperature before embedding in paraffin for histological analysis. Instruments were disinfected between samples by wiping off grossly visible tissue, then placing them in 15% bleach for 10 min, followed by ethanol for another 5 min, and then rinsed in distilled water.

Histopathology was conducted on all major tissues using standard hematoxylin and eosin stain. Immunohistochemistry for West Nile virus antigen was performed using validated protocols on selected tissues (data not shown).

### Diagnostic Testing

All diagnostic testing was performed at the Pennsylvania Animal Diagnostic Laboratory System - New Bolton Center laboratories, University of Pennsylvania, School of Veterinary Medicine, Kennett Square, PA. Toxicology screens included ACR rodenticides, heavy metals and organic chemicals (data not shown). Targeted PCR were used to test for the presence of Salmonella, avian influenza A virus, avian paramyxovirus type 1 (Newcastle Disease), and Chlamydia (data not shown). All standard methods were developed and validated either in-house or by the U.S. Department of Agriculture’s National Veterinary Services Laboratory in Ames, Iowa.

### mNGS library preparation

Frozen tissue samples from the brain, liver, lung, and spleen of three representative cases found deceased on one of the eastern Pennsylvania sites, along with similar tissue samples from three control vultures that died from noninfectious causes at the same site in 2022 and 2023, were processed for metagenomic sequencing. Cases were arbitrarily labeled Bird 1, 2 and 3 for ease of discussion and comparison. Tissue samples were aliquoted into bead-bashing tubes (ZR Bashing Bead Lysis Tubes S6012-50, 0.1 & 0.5 mm, Zymo Research) with at least 400 µL of DNA/RNA Shield (Zymo Research), homogenized at 30 Hz for 1 min, incubated on ice for 1 min, and homogenized again for 1 min at 30 Hz. 200 µL of bead-bashed sample supernatant in 1X Shield was used for total nucleic acid extraction utilizing the quick-DNA/RNA Pathogen Magbead kit (Zymo Research). RNA was extracted using DNAse according to the manufacturer’s protocol and quality checked on the TapeStation 4200 (Agilent Technologies) for RNA integrity using the High Sensitivity RNA ScreenTape (Agilent Technologies). ERCC spike-in (RNA standard dilution series from External RNA Controls Consortium) was used as a positive control and water as a negative control for background contamination. FastSelect - rRNA HMR (Qiagen) was used for mammalian RNA ribosomal depletion at 1:10X. NEBNext Ultra II Library Prep kit (New England Biolabs) was used to reverse transcribe RNA into cDNA which was used to construct and barcode sequencing libraries. The RNA sequencing libraries underwent 150-nt paired-end Illumina sequencing on a NextSeq2000 Sequencing System (Illumina) with a target of at least 5 million reads per sample.

### mNGS bioinformatic analysis using CZ ID

CZ ID, an open-source sequencing analysis platform which identifies microbes in mNGS data, was used for microbe and pathogen detection (http://czid.org<u>, v8.0</u>). The pipeline trims adapters, low-quality, and low-complexity reads using fastp ^18^, filters out human and vulture host reads with Bowtie2 ^19^ and HISAT2 ^20^, then removes duplicated reads with CZID-dedup (https://github.com/chanzuckerberg/czid-dedup). Remaining reads are queried against the NCBI nucleotide (NT) and non-redundant protein (NR) database using GSNAP ^21^and RAPSearch2 ^22^, respectively, to identify and determine microbial taxa ^23^.

Water control samples were used to account for background contamination by creating a mass-normalized background model for tissue samples. Significant microbial hits were called using the following threshold filters mapping to specific genera or species normalized by reads per million (rPM); z-score ≥ 1 (to denote significant presence of microbe in sample compared to water background), NT rPM ≥ 10 (at least 10 reads per million aligned at the nucleotide level to specific taxa), NR rPM ≥ 5 (at least 5 reads per million aligned at the amino acid level to specific taxa), and average NT alignment length ≥ 50 base pairs. To further increase validity and reliability of the microbial hits, both NT and NR sequence filters were employed to minimize potential false positive detection which could have resulted from low quality alignments to the NCBI database.

### Genome recovery and characterization

Reads mapping to the *Bandavirus* genus were assembled into contigs in CZ ID with SPAdes ^24^ and were downloaded and further explored using Geneious Prime (2024.0.7). Bandavirus reads identified in CZID were mapped to contigs and visually inspected. Regions with coverage < 10 reads or > 10 reads with signs of read misalignment were trimmed from the contigs. A simple majority rule was applied for base calls for positions within contigs that had mixed base calls.

Multiple sequence alignments of the recovered contigs were performed with MAFFT ^25^ to compare the sequence similarity across each tissue sample and bird. BlastX was used to identify closest reference sequences for each contig genomic segment. The top reference sequence was Hunter Island virus (GenBank accession numbers: <u>NC_027717</u>, <u>NC_027716</u>, <u>NC_027715</u>). Geneious software was used to identify open reading frames within each of the recovered genome segments, and to compare their size and sequence similarity to counterparts in the Hunter island virus. PySimPlot (https://github.com/jonathanrd/PySimPlot; Lole et al. 1999) was used to perform scanning pairwise nucleotide identity plots of each segment and scanning pairwise amino acid identity plots of the corresponding ORFs within each segment, using Hunter Island virus as the reference sequence. A window size of 50 incremented by 1 unit was used for all the pairwise identity comparisons. Results were visualized with GraphPad Prism (v9.5.1).

### Phylogenetic analyses

Four maximum likelihood phylogenies for each segment and the corresponding proteins they encoded (i.e. polymerase, glycoprotein, nucleoprotein and nonstructural protein) were constructed to examine the evolutionary relatedness of the recovered viral segments to other viral genera in the *Phenuiviridae* family. A representative set of viral genomes and their respective coding sequences were used for this (summarized in **Supplementary Table 2**).

MAFFT ^25^was used to generate multiple sequence alignments (MSA) of the translated proteins, and ModelTest-ng was utilized to determine the best-fit amino acid substitution model for each protein MSA ^27,28^. Phylogenies were constructed with IQTREE ^29^ and figures were viewed and generated from treefile on FigTree v1.4.4 (http://tree.bio.ed.ac.uk/software/figtree/) and Figma.

### RT-qPCR diagnostic

The sequences of the L genome segment recovered from Bird 2 were used as input for the RT-qPCR primer design for virus detection, and the region that encodes the RdRp protein was targeted. Primers and probes (forward primer = 5’-GGCTCATGGGTGAGGTATT-3’, reverse primer = 5’-GGGAGGTTCACTAGATTGGTTAG-3’, probe = 5’-ACACAATAACCTCAGGAGACTGGGAGA-3’) were generated with the IDT PrimerQuest tool (https://www.idtdna.com/PrimerQuest/Home/Index).

RT-qPCR testing of RNA extracted from spleen or liver samples collected from suspected cases were prioritized for testing, since these tissues yielded the highest number of viral reads in the mNGS analysis. Parallel tissues collected from a single black vulture that died in 2023, under circumstances unassociated with the mortality event, were included as controls for the testing. Tissues were typically provided frozen and thawed before dissecting. Approximately 100 mg of tissue were harvested using a sterile scalpel, then placed in a Precellys 2 mL Hard Tissue Homogenizing Ceramic Beads CK28 tubes (Bertin Technologies) containing 1 mL of 1X PBS. Tissues were next homogenized for 8 cycles of 30 s at 18 Hz in a Precellys Evolution Touch Homogenizer (Bertin Technologies). After homogenization, the tubes were centrifuged at 1500 *x g* for 3 min, followed by an additional step of 3000 *x g* for 3 min to ensure proper separation of tissue debris from the supernatant. 600 µL of supernatant was transferred into clean labeled microcentrifuge tubes. RNA extractions were performed on 140 µL of this material with the QIAamp Viral RNA Mini Kit (QIAGEN), following the manufacturer’s instructions. Samples were eluted in 60 µL, and 7 µL of this RNA prep was used as input for RT-qPCR diagnostic with the Luna Universal One-Step RT-qPCR kit (New England Biolabs) in a QuantStudio 5 Real-time PCR thermocycler (ThermoFisher Scientific) for 45 PCR cycles.

## Data availability

Near-complete bandavirus genome segment sequences recovered in this study were deposited to GenBank (**Supplementary Table 1**, Accessions PQ442242-PQ442257). The raw sequence data can be accessed via SRA BioProject PRJNA1198986 (biosample accessions SAMN45838115 - SAMN45838138; fastq accessions SRR31744714 - SRR31744737).

## Results

### Initial necropsy and histopathology

The majority of the 27 birds examined at New Bolton Center were females (n=23) and presented in very good body condition, suggestive of acute death. Data on age of the affected birds were incomplete however, of those recorded 16/22 were adults and 6/22 were noted as immature. Gross examination of internal organs revealed enlarged livers characterized by pale yellow streaks and enlarged spleens **(Figure 1A)**, with evidence of hemorrhage in the body cavity and occasional visceral gout **(Figure 1B)**. Multiple liver and/or splenic lesions were noted in 23/27 birds, while 4/27 suspected cases had no gross abnormalities of the liver and spleen, or signs of hemorrhage **(Table 1**). Necropsy results of one of the birds (PA-22) sent to an independent diagnostic lab were reported back to New Bolton Center and are also included in **Table 1**.

**Figure 1.**
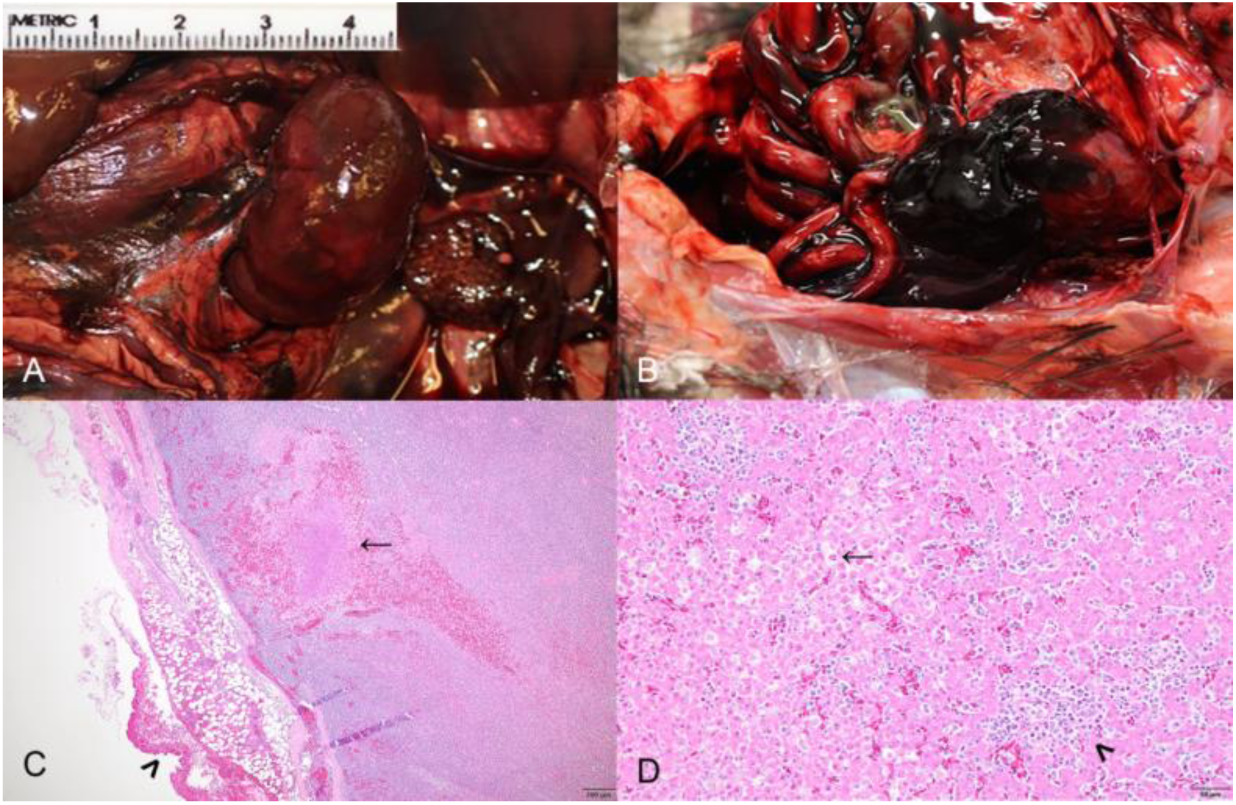
Gross pathology and histological analysis of splenic and liver lesions. Gross (**A and B**) and microscopic (**C and D**) lesions of bandavirus infection in black vultures. **A**, A grossly enlarged and congested spleen in an infected vulture. **B**, An infected vulture with splenic rupture with partially clotted blood coating the capsular surface and spilling out into the surrounding coelom. **C**, Spleen from an infected vulture showing foci of fibrin, necrosis (arrow), and hemorrhage within the parenchyma as well as capsular hemorrhage on the splenic surface (arrowhead); hematoxylin and eosin (HE) stain. **D**, Liver from an infected vulture with individual hepatocellular necrosis (arrow) and mild sinusoidal expansion by mixed round cells (arrowhead); HE stain.

**Table 1.**
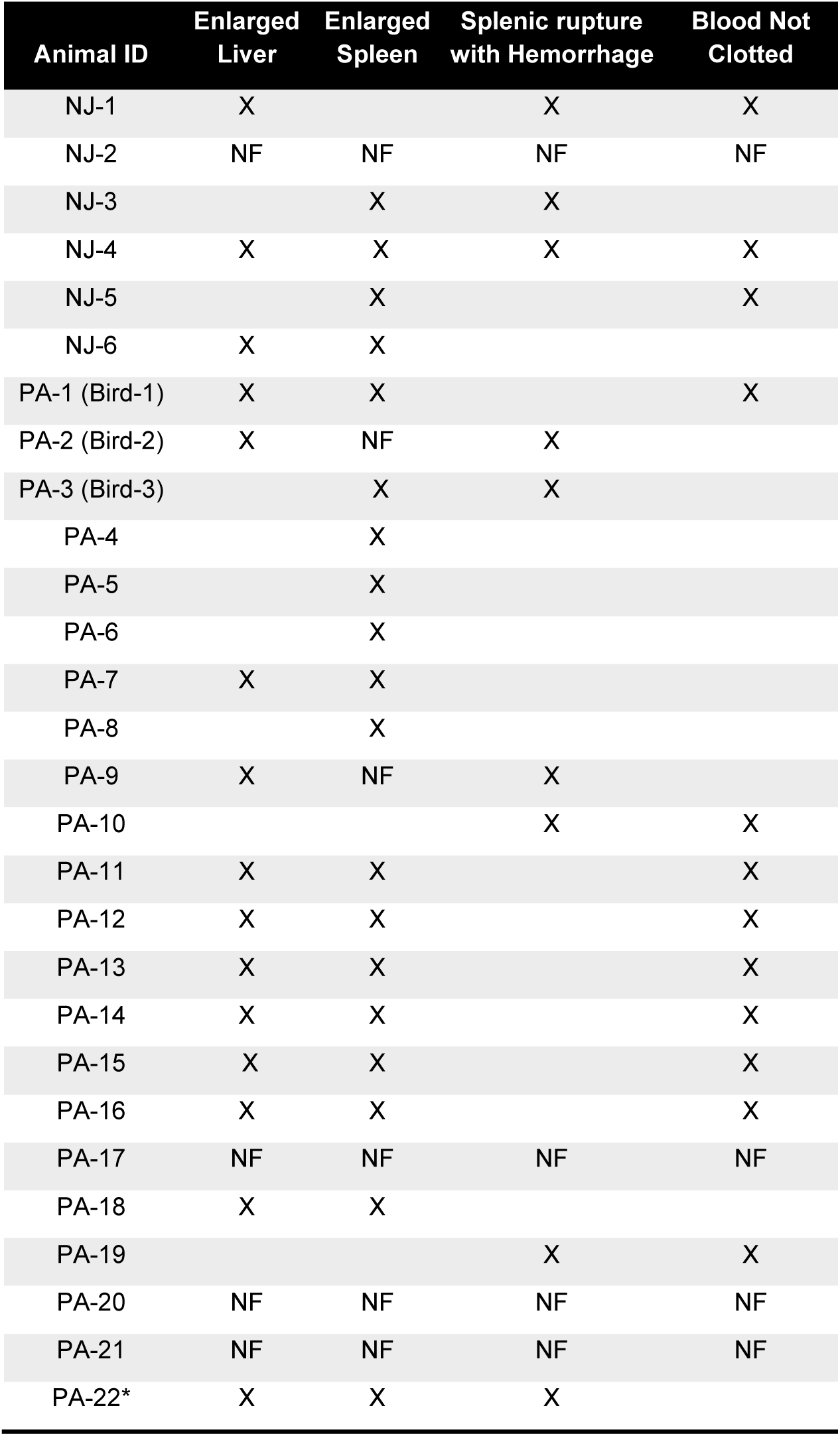
Gross pathology results from suspected cases. NF indicates “no findings noted” upon examination. *PA-22: Necropsy results reported back to New Bolton Center from external site.

Histopathological examination of the splenic lesions revealed congestion, hemorrhage, and/or necrosis/infarcts that were occasionally associated with splenic rupture and/or coelomic hemorrhage **(Figure 1C)**. Both livers and spleens exhibited significant lymphocytic infiltrate **(Figure 1D)**. Toxicology revealed rodenticide exposure, but at low concentrations, consistent with chronic exposure from multiple dietary sources rather than acute toxicity, and was thus ruled out as a likely primary cause of mortality (data not shown). Testing for aerobic and anaerobic bacterial agents revealed the presence of commensal, environmental and potentially opportunistic bacteria (data not shown), but nothing that was in common across all cases.

Specific PCR testing for Salmonella, Chlamydia, avian influenza A virus, avian paramyxovirus type 1 (virulent Newcastle Disease), and immunohistochemistry for West Nile Virus were all negative, along with limited virus isolation testing (data not shown). Since 2020, there have been no reports of additional large mortality events among vultures in these geographic areas, and in the absence of any clear causal findings via conventional diagnostic approaches, the mortality event remained unresolved.

### mNGS analysis for pathogen detection

In 2023, mNGS analysis was performed to interrogate tissues from three representative cases and three control vultures, and successfully identified distinct microbial sequences present in the case birds. Reads aligning to the *Phlebovirus* and *Bandavirus* genera were detected across all tissues collected from the case birds, and none of the tissues collected from the control birds. Quantitative taxonomic analysis of the recovered viral reads revealed that the majority of the reads in the case birds aligned to sequences assigned to the bandavirus genus **(Figure 2A).**

**Figure 2.**
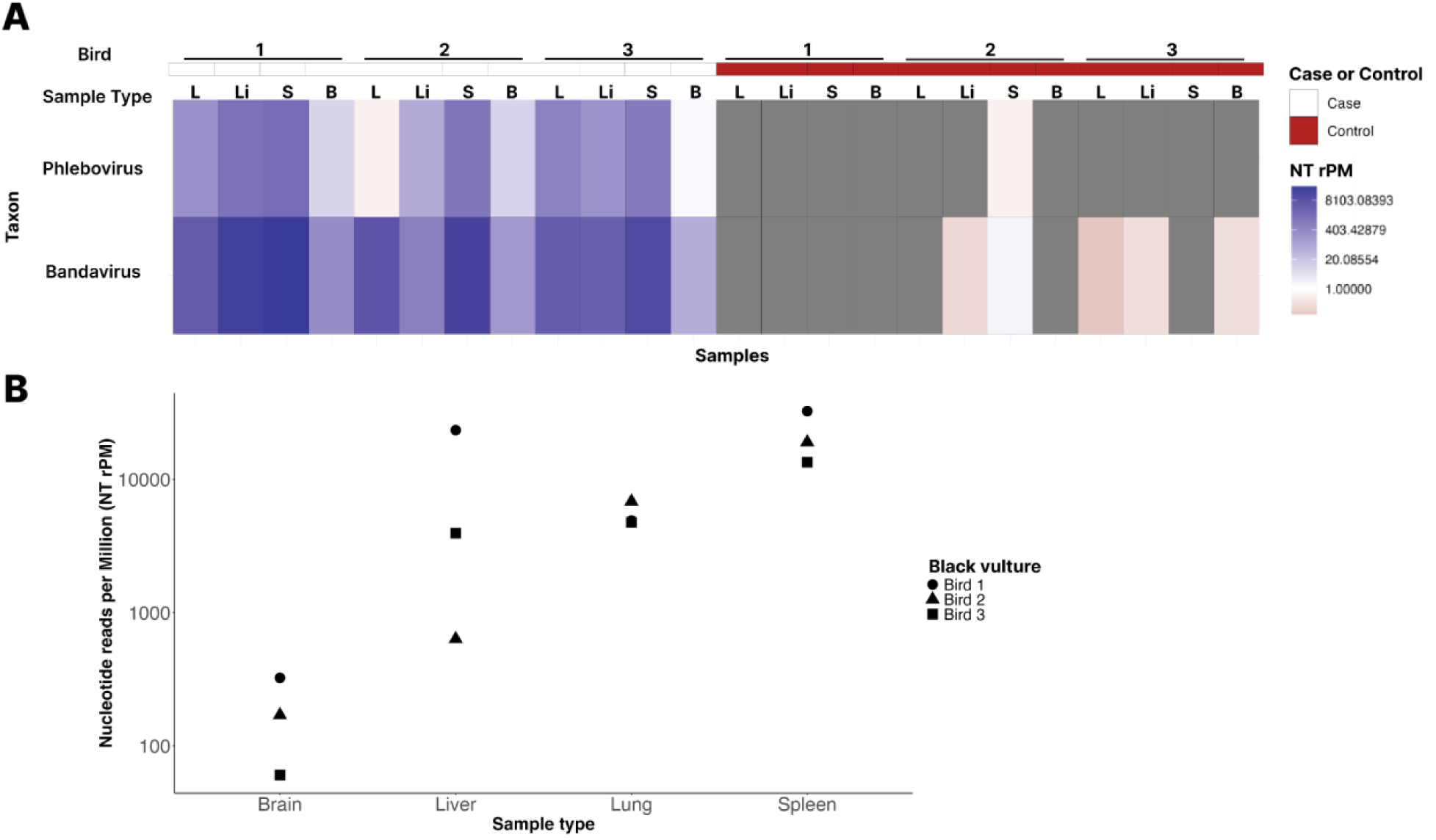
Bandavirus detection across all sample types of black vulture cases. **A**, Microbe detection of all viruses within the black vultures at the genus level for each respective bird; the color denotes if it was a case (white) or control (red); the sample type key: L - Lung, Li - Liver, S - Spleen and B - Brain. **B**, Genus-level bandavirus detection across all the case sample types (x-axis) with their relative abundance NT rPM (y-axis). The shape denotes the bird.

Overall, a significant fraction of the non-host reads recovered from each of the case birds corresponded to bandavirus reads: 56% in Bird 1, 16% in Bird 2 and 12% in Bird 3 **(Supplementary Table 1).**

Quantification of the bandavirus reads recovered within the tissues of the case birds revealed different patterns of bandavirus read abundance; however, in all case birds, we consistently observed the highest recovery of bandavirus reads from splenic tissue **(Figure 2B, Supplementary Table 1)**. Likewise, for all tissues of all the case birds, we observed greater than 10-fold more bandavirus reads recovered by alignment to the NCBI non-redundant (NR) protein database compared to the nucleotide (NT) database. Moreover, the recovered NR bandavirus reads were readily assembled into contigs of near-complete genome segment lengths **(Supplementary Table 1)**. This bias in recovery of bandavirus reads via NR alignment compared to NT alignment is consistent with the presence of a potentially novel bandavirus isolate, with a distinct nucleotide sequence that encodes a protein sequence similar to previously described bandaviruses.

The recovery of bandavirus contigs across multiple tissues within each case bird allowed us to determine if a single or multiple bandaviruses were associated with the mortality event. We performed multiple sequence alignment of the complete protein coding regions of each viral genome segment to understand the bandavirus sequence similarity both within and between the case birds. Within Bird 2, near-complete bandavirus genome length L, M and S genome segments were recovered from multiple tissues **(Table 2, Supplementary Table 1)**. The amino acid and underlying nucleotide sequences for the protein-coding regions of each of these segments were identical. Within Bird 1 and Bird 3, contigs corresponding to only a subset of the genome were recovered from multiple tissues. For example, in Bird 3, near-complete bandavirus genome length M and S segments could be analyzed, while in Bird 1, it was only feasible to examine the S genome segment. We found the amino acid and underlying nucleotide sequences of each segment that could be analyzed within both Bird 1 and Bird 3 were identical. Taken together, these data indicate that a single bandavirus isolate was present within each of the case birds.

**Table 2.** Amino acid identity of the recovered bandavirus and other bandavirus reference genomes to the Hunter Island virus reference genome.

| Segment | Reference viral species | % Identical sites to Hunter Island virus |
| --- | --- | --- |
| L | Recovered bandavirus | 85.3 |
|  | Heartland virus | 66.8 |
|  | SFTS | 65.5 |
|  | SFTS 2 | 65.7 |
|  | Guertu virus | 66.5 |
| M | Recovered bandavirus | 75.2 |
|  | Heartland virus | 53.7 |
|  | SFTS | 54.3 |
|  | SFTS 2 | 54.4 |
|  | Guertu virus | 53.2 |
| S nonstructural protein | Recovered bandavirus | 68.5 |
|  | Heartland virus | 27.8 |
|  | SFTS | 17.6 |
|  | SFTS 2 | 32.3 |
|  | Guertu virus | 23.5 |
| S nucleoprotein | Recovered bandavirus | 90.8 |
|  | Heartland virus | 28.6 |
|  | SFTS | 0.4 |
|  | SFTS 2 | 61.5 |
|  | Guertu virus | 20.0 |

We next performed multiple sequence alignment of representative viral genome segments recovered from each bird to examine if the same bandavirus isolate was recovered in all 3 of the case birds. For the L genome segment analysis, contigs from only Bird 1 and Bird 2 sequences contributed to the analysis. In contrast, for the M and S genome segments, it was possible to compare contigs recovered from all 3 birds. For all of these between-bird genome segment comparisons, we found both the amino acid sequences and underlying nucleotide sequences were identical, indicating infection with a single bandavirus isolate was associated with this mortality event.

### Bandavirus sequence analysis and phylogenetics

To shed light on the relationship of this potentially novel bandavirus recovered from cases of the mortality event to other previously described bandaviruses, we performed blastX alignment of the assembled bandavirus contigs. The best matches corresponded to 3 different bandaviruses: Hunter Island virus (*Bandavirus albatrossense)*, Heartland virus, and SFTSV. Among these, the most closely related virus was Hunter Island virus which showed the highest amino acid percentage identity across all segments of the recovered virus, and was selected to be the reference genome for analyses. We next performed scanning pairwise sequence identity analysis comparing the recovered virus with the Hunter Island reference at both the nucleotide and amino acid levels to gain a higher resolution view **(Figure 3).** We selected a representative set of recovered contigs with lengths similar to the reference genome for this analysis. We first examined the L genome segment, which encodes the bandavirus RdRp, as this sequence is used for taxonomic characterization. An RdRp coding region with less than 90% amino acid identity is indicative of a novel *Bandavirus* species as outlined by the ICTV guidelines. The representative L bandavirus segment used for this analysis was recovered from the lung of Bird 1. It has a length of 6353 nt, 5 nt shorter than the reference with a length of 6358 nt. This contig contains an RdRp open reading frame with a length identical to the Hunter Island Virus RdRp sequence (6258 nt), and showed an average percent identity of 74.9% (range: 34-92%) at the nucleotide level and 85.3% (range: 52-100%) at the amino acid level to the Hunter Island virus RdRp sequence **(Figure 3A)**. The representative M segment we used for this analysis was derived from the liver of Bird 2, with a length of 3309 nt, is 19 nt shorter than the Hunter Island Virus M reference (3328 nt). The recovered M segment harbors a 3189 nt open reading frame encoding the bandavirus glycoprotein. This is shorter than the Hunter Island Virus glycoprotein coding sequence by 3 nt. Moreover, a deletion of a single amino acid (position 529 in Hunter Island virus glycoprotein) was observed in this open reading frame and all the glycoprotein open reading frames recovered from the M segment contigs in this study. Aside from this difference, the scanning pairwise sequence analysis revealed an overall average nucleotide sequence identity of 68.3% (range: 42-90%) and amino acid identity of 75.2% (range: 44-100%) to the Hunter Island virus M segment sequences **(Figure 3B)**. The representative bandavirus S segment was derived from Bird 3, with a length of 1682 nt, one nucleotide longer than the reference sequence. Similar to other bandaviruses, the recovered S segment shows a characteristic ambisense coding potential with a 65 nt intergenic region to allow for each protein to be translated independently ^30^. The nucleoprotein open reading frame was identical to the reference of 747 nt (248 aa), as was the nonstructural protein open reading frame at 804 nt (267 aa). The recovered bandavirus S segment showed an overall 71.8% average nucleotide sequence identity (range: 32-94%) to the Hunter Island virus S segment, and 90.8% (range: 84- 96%) and 68.5% (range: 40-84%) amino acid identity to the Hunter Island virus nucleoprotein and nonstructural protein sequences, respectively. **(Figure 3C)**.

**Figure 3.**
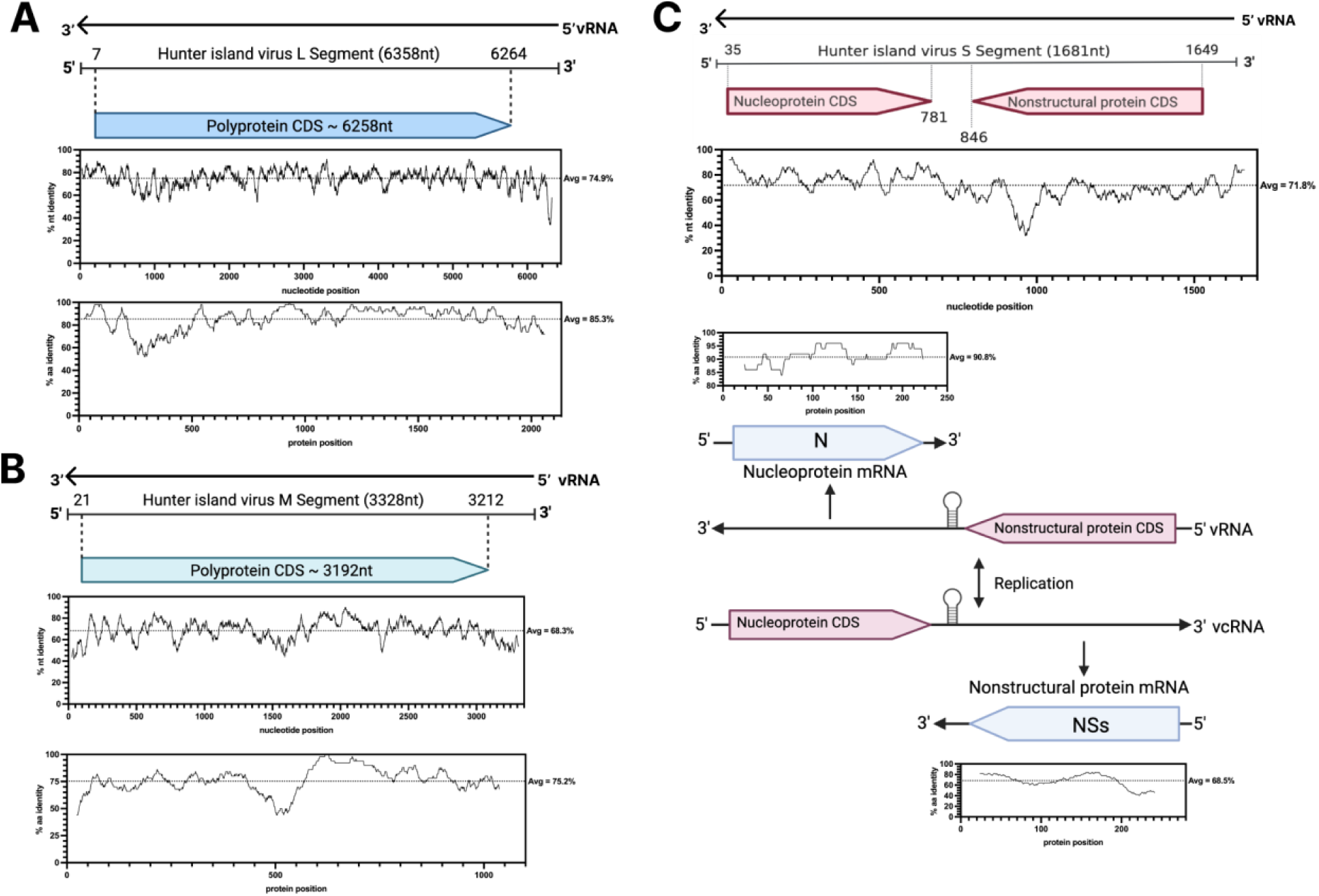
Recovered bandavirus sequence similarity to Hunter Island virus. **A,** Hunter Island Virus reference genome L segment (GenBank accession <u>NC027717</u>) annotated with the coding region of the polymerase gene, and scanning nucleotide and amino acid pairwise identity plots of a representative L segment contig of the recovered bandavirus (GenBank accession PQ442249). **B,** Hunter Island virus reference genome M segment (GenBank accession <u>NC027715</u>) annotated with the coding region of the glycoprotein gene, and scanning nucleotide and amino acid pairwise identity plots of a representative M segment contig of the recovered bandavirus (GenBank accession PQ442256). **C,** Hunter Island virus reference genome S segment (GenBank accession <u>NC027716</u>) annotated with the coding regions of the nucleoprotein and nonstructural protein genes, and the scanning nucleotide and amino acid pairwise identity plots of a representative S segment contig of the recovered bandavirus (GenBank accession PQ442248).

The level of amino acid sequence identity between the recovered bandavirus and Hunter Island virus is distinctly higher than that observed for other bandavirus species **(Table 2)**, indicating that the newly recovered virus is the most closely related bandavirus to Hunter Island virus. To assess where the recovered bandavirus fits within the *Phenuiviridae* viral family, we performed a phylogenetic analysis of the protein sequences encoded by each segment. For the RdRp and the glycoprotein phylogenies, the bandaviruses cluster together with the recovered virus mapping closest to the Hunter Island virus reference **(Figure 4A and B, respectively)**. For the nucleoprotein and nonstructural protein phylogenetic analyses, the bandaviruses split into three **(Figure 4C)** or two distinct clusters **(Figure 4D)**, respectively; with the recovered bandavirus consistently mapping closest to the Hunter Island virus reference **(Figure 4C and D)**.

**Figure 4.**
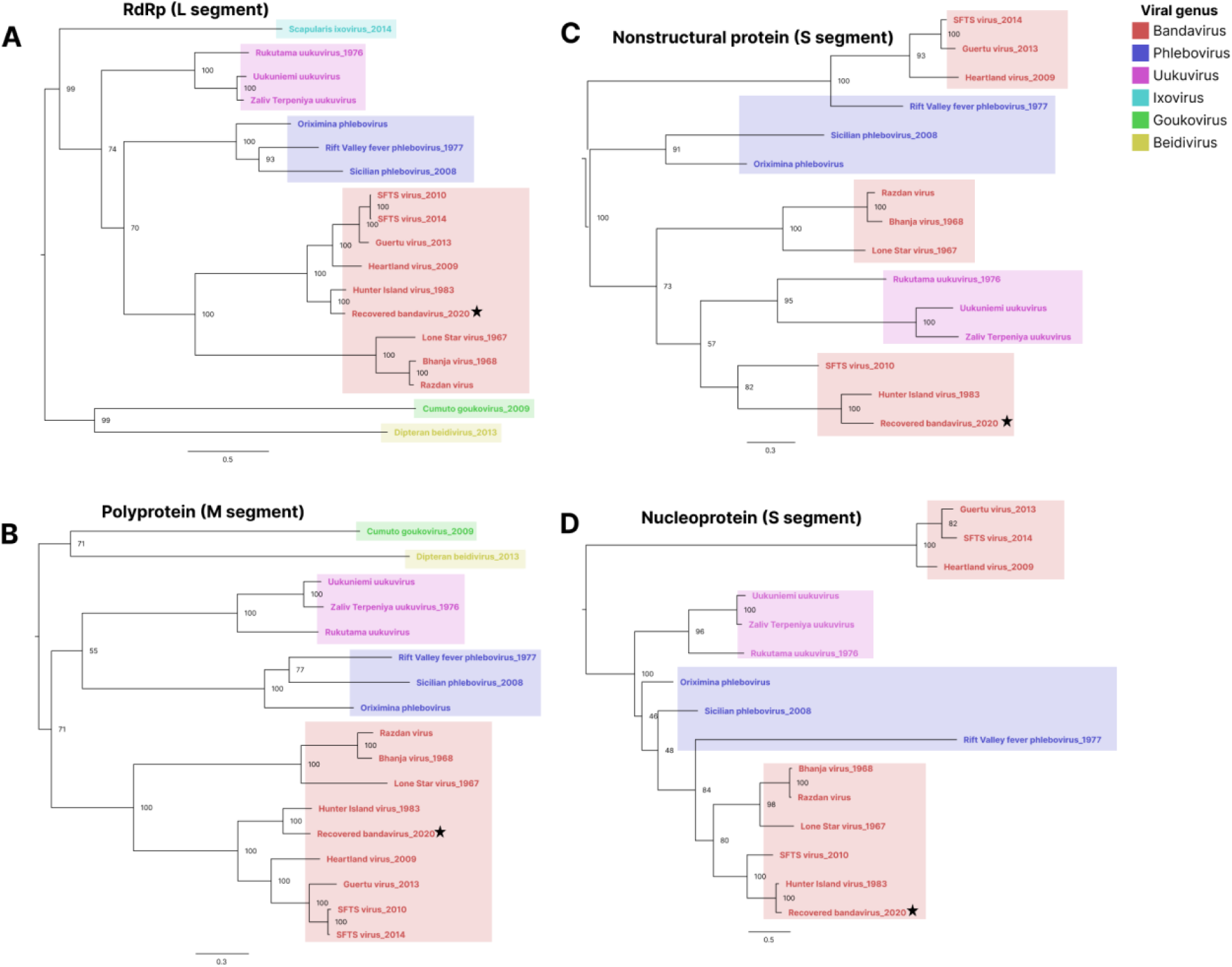
Phylogenetic analysis of representative *Phenuiviridae* genus proteins with the recovered genome. Phylogenetic trees based on the amino acid sequences of the L segment RNA-dependent polymerase **(A),** the M segment polyprotein (glycoprotein) **(B)**, and the S segment nonstructural protein **(C)** and nucleoprotein **(D)**. All trees are midpoint rooted. In all panels, colors denote the viral genus and the year denotes when the respective viruses were identified. The star indicates the genome recovered in this study. Scale bar represents amino acids substitutions per site.

### Targeted RT-qPCR screening for bandavirus

To investigate the association between infections with the recovered bandavirus and the fatalities associated with the 2020 die off event, we developed primers and probes to enable targeted RT-qPCR screening of the remaining suspected cases for infection with the recovered bandavirus. The results from this assay confirmed the presence of the novel bandavirus in all of the suspected cases with an average C_t_ value of 25 (range 17-35) **(Figure 5, Supplementary Table 3)**. The negative control exhibited undetermined Ct value above 40.

**Figure 5.**
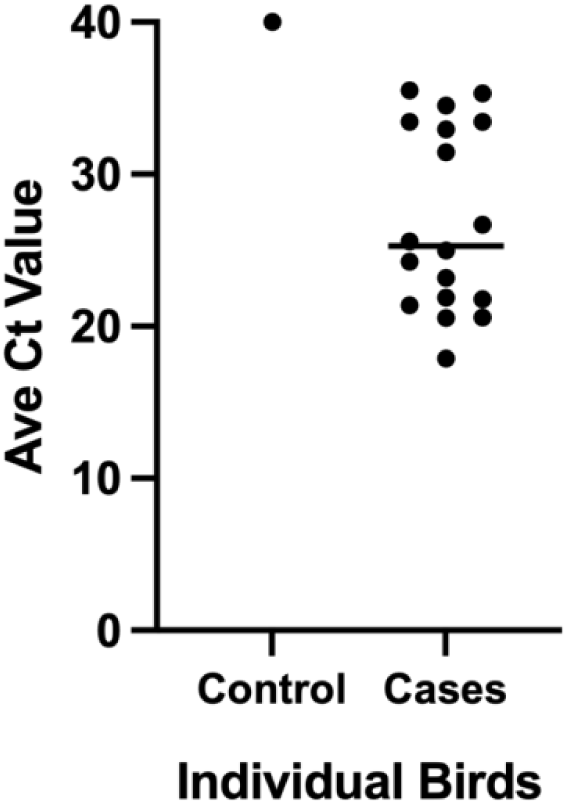
Cycle threshold (C_t_) values obtained during RT-qPCR in control and cases. Targeted PCR was performed on eighteen of the suspected cases. A healthy black vulture that died of trauma, unrelated to the mortality event, was used as a negative control. Average C_t_ values are represented, and values <40 were designated as positive for the bandavirus. Actual C_t_ values for each bird are listed in Supplementary Table 3.

## Discussion

Here, we applied mNGS to investigate the cause of a black vulture die-off that occurred in eastern Pennsylvania and western New Jersey over the course of 5 months in 2020. We leveraged this unbiased approach in the context where conventional testing failed to identify pathogens, to obtain a comprehensive overview of all the microbes within the samples available from the birds collected for analysis. We examined tissue samples from three of the deceased vultures and detected a new bandavirus. These results were extended to all the suspected cases using targeted RT-qPCR. Taken together, along with the known natural history of related viruses, these data strongly suggest that the causative agent of this mortality event was a novel species of bandavirus. We propose to name this additional member *Bandavirus coragypsense* after the black vulture genus name *Coragyps*.

Some bandavirus species have been reported to cause outbreaks involving avian, human and other mammalian hosts ^31^. Symptoms associated with bandavirus infections in humans range from fever, vomiting, diarrhea, headache and thrombocytopenia ^32,33^. Clinical manifestation of thrombocytopenia includes enlarged spleen caused by trapping of platelets in the spleen ^34^.

Notably, splenomegaly was also observed in the deceased birds. Virulence across bandavirus species that infect humans can vary considerably. For example, Heartland virus has been reported to cause hospitalization of acutely infected individuals that eventually recover ^35–38^, while Dabie bandavirus, the source of the largest human *Bandavirus* outbreaks across Asia, has a reported mortality rate ranging from 10-30% of those infected ^30^. Thus, it is important to understand the phylogeny of novel members of this genus in order to gain insight into their biology and potential for zoonotic spillover.

Sequence and phylogenetic analysis showed that the recovered novel bandavirus is most closely related to Hunter Island virus. Hunter Island virus was discovered in 1983 during a large die-off of young Shy albatrosses (*Thalassarche cauta*) on Albatross Island off the coast of Tasmania, Australia. In that event, 50% of the affected birds tested positive for the virus ^39^. In 2002, another albatross disease outbreak was reported on the same island and the virus was detected in tick (*Ixodes eudyptidis*) pools found on diseased and healthy birds. However, in this event the virus was not detected in bird serum ^40^. The phylogenetic groupings of genomic segments from the recovered bandavirus and Hunter Island virus suggest they are closely related, but likely to be distinct species as indicated by their low L segment RdRp coding region sequence similarity.

Ticks have been previously identified as a common vector for bandavirus transmission, as various viral species have been isolated from ticks across five continents ^31^. Anecdotal evidence has associated the presence of a tick bite prior to disease onset in some human cases and more recently, Heartland virus was isolated from the Lone Star tick *Amblyomma americanum* in Illinois and New York, implicating this species as a likely vector of infection ^33,41,42^. Furthermore, the distribution of Dabie bandavirus throughout East Asia is characteristic of a tick and migratory bird pattern of spread, highlighting an important vector-reservoir dynamic for the geographical reach of these viruses ^30^. While ticks are thought to be the primary vector, cases of person-to-person spread have also been reported after exposure to blood or body fluids of severely ill patients ^32^.

As this was a retrospective study we were not able to look for ticks in the nests of the affected vultures, nor did we find any ticks on the deceased vultures, and therefore could not determine if a tick species was also the source of infection in this outbreak. Screening ticks, especially newer species, in the regions where the die-off occurred would aid in determining the possibility of a tick vector. Presence of the virus in ticks, however, does not preclude potential bird-to-bird spread of the infection as an epidemiological mechanism. Moreover, since vultures feed on carrion, they could also have been infected by an intermediate host that was harboring the virus, thus warranting the sampling of other fauna in the region for prior exposure to this virus as well. In addition to helping understand transmission dynamics of the virus, testing of migratory birds that travel along the Northeastern U.S. may provide insight into how this virus was first introduced into the black vulture colonies that reside there. Vultures could have been exposed to the virus via direct bird-bird interaction with transient birds or via the introduction of infected tick species carried in from other locations.

In summary, we employed mNGS to successfully identify a new bandavirus species as the likely cause of a black vulture mortality event, underscoring the importance of employing novel methods when routine testing cannot provide answers. As a retrospective study, our investigation was limited to sampling the deceased birds, and thus the source of this virus remains unclear and an area for further investigation. The development of simple molecular tests, such as the RT-qPCR assay described here, can be easily employed for future investigations focused on identifying potential vectors, such as invasive tick populations that have been associated with the introduction of other bandaviruses ^43^. In addition, serosurveillance of other local fauna and of migratory birds can help elucidate the overall prevalence of exposure and point to potential viral reservoirs. Bandaviruses, as well as other members of the *Phenuiviridae* family, have been shown to be important sources of morbidity and mortality in humans as a result of zoonotic spillover. Our investigation emphasizes the need for more coordinated surveillance and real-time response to animal mortality events to identify important causes of disease. These results also highlight the utility of employing metagenomic sequencing to discover new sources of previously unknown zoonoses with epidemic potential, better preparing us for what may lie ahead.

## Acknowledgements

AK and CMT are supported by grants from Chan Zuckerberg Initiative; SJB, SD, KDN, MLG and LAM are supported by 500361: Diagnose Disease of Animals and Coordinate Diagnostic Efforts with Other Participants of the PA Animal Diagnostic System; EAM, KDN, MLG, RBG, LAM are supported by 501097: Comprehensive Wildlife Health Program for the State of Pennsylvania; AOGH is supported by 584822: Stewardship of Wildlife Health Initiative.

