## Supplemental Data for "Discovery of a novel bandavirus using metagenomic sequencing in a retrospective analysis of an unresolved 2020 mortality event involving black vultures in the northeastern United States"

### Supplementary Table Headings

**Supplementary Table 1.** Summary of sample metadata, mNGS results, bandavirus read and contig yields.

**Supplementary Table 2.** Reference genomes used for pairwise sequence analyses summarized in Table 2, and phylogenetic analysis in Figure 4.

**Supplementary Table 3.** PCR Cycle threshold ($C_t$) values by animal ID for each bird tested and represented in Figure 5. NA denotes samples that were “not available” for bandavirus RT-qPCR testing.

### Supplementary Tables

| Library_ID | SRA_Biosample_Accession | SRA_Accession | Animal_ID | Bird_ID | Case or Control | sample_type | total_reads | passed_filter_reads | NT_bandavirus_reads | NR_bandavirus_reads | NR_bandavirus_reads/NT_bandavirus_reads | Tissue_Percent_BandavirusReads/PassedFilters | WholeBird_Percent_BandavirusReads/PassedFilter | GenbankAccessionNumbers |
|---|---|---|---|---|---|---|---|---|---|---|---|---|---|---|
| RR167_066_Lung_S71 | SAMN45838115 | SRR31744737 | N2017664 | 1 | Case | Lung | 3240208 | 475892 | 15205 | 204632 | 13 | 43 | 56 | PQ442249 |
| RR167_067_Liver_S72 | pending | pending | N2017664 | 1 | Case | Liver | 9785562 | 4325980 | 183015 | 2,347,210 | 13 | 54 | | |
| RR167_068_Spleen_S73 | SAMN45838117 | SRR31744736 | N2017664 | 1 | Case | Spleen | 7076948 | 3986114 | 224591 | 3181494 | 14 | 80 | | PQ442252 |
| RR167_069_Brain_S74 | SAMN45838116 | SRR31744725 | N2017664 | 1 | Case | Brain | 5866906 | 1536312 | 1876 | 19797 | 11 | 1 | | PQ442250, PQ442251 |
| RR167_070_Lung_S75 | SAMN45838118 | SRR31744720 | N2017665 | 2 | Case | Lung | 12799086 | 5742880 | 55342 | 876672 | 16 | 15 | 16 | PQ442253 |
| RR167_071_Liver_S76 | SAMN45838120 | SRR31744719 | N2017665 | 2 | Case | Liver | 9415792 | 6965530 | 5012 | 140272 | 28 | 2 | | PQ442242, PQ442256 |
| RR167_072_Spleen_S77 | SAMN45838121 | SRR31744718 | N2017665 | 2 | Case | Spleen | 8250984 | 4448242 | 130178 | 2158324 | 17 | 49 | | PQ442243 |
| RR167_073_Brain_S78 | SAMN45838119 | SRR31744717 | N2017665 | 2 | Case | Brain | 6540852 | 2772114 | 1098 | 16249 | 15 | 1 | | PQ442254, PQ442255, PQ442257 |
| RR167_074_Lung_S79 | SAMN45838123 | SRR31744716 | N2017668 | 3 | Case | Lung | 6284250 | 5287666 | 29343 | 817951 | 28 | 15 | 12 | PQ442245, PQ442247 |
| RR167_075_Liver_S80 | SAMN45838122 | SRR31744715 | N2017668 | 3 | Case | Liver | 17153800 | 8820168 | 38071 | 503415 | 13 | 6 | | PQ442244 |
| RR167_076_Spleen_S81 | SAMN45838125 | SRR31744714 | N2017668 | 3 | Case | Spleen | 5453852 | 4373800 | 72244 | 1229783 | 17 | 28 | | PQ442248 |
| RR167_077_Brain_S82 | SAMN45838124 | SRR31744735 | N2017668 | 3 | Case | Brain | 11838404 | 3370180 | 704 | 71322 | 101 | 2 | | PQ442246 |
| RR167_078_Lung_S83 | SAMN45838130 | SRR31744734 | N2302788 | 4 | Control | Lung | 23611568 | 4576254 | 0 | 0 | 0 | 0 | 0 | |
| RR167_079_Liver_S84 | SAMN45838131 | SRR31744733 | N2302788 | 4 | Control | Liver | 8929080 | 6885188 | 0 | 0 | 0 | 0 | | |
| RR167_080_Spleen_S85 | SAMN45838132 | SRR31744732 | N2302788 | 4 | Control | Spleen | 357050 | 3458 | 0 | 15 | NA | 0 | | |
| RR167_081_Brain_S86 | SAMN45838133 | SRR31744731 | N2302788 | 4 | Control | Brain | 10629554 | 6113508 | 0 | 2 | NA | 0 | | |
| RR167_082_Lung_S87 | SAMN45838134 | SRR31744730 | N2302789 | 5 | Control | Lung | 848998 | 261860 | 0 | 14 | NA | 0 | 0 | |
| RR167_083_Liver_S88 | SAMN45838135 | SRR31744729 | N2302789 | 5 | Control | Liver | 6774462 | 2273546 | 1 | 16 | 16 | 0 | | |
| RR167_084_Spleen_S89 | SAMN45838136 | SRR31744728 | N2302789 | 5 | Control | Spleen | 2107046 | 558138 | 3 | 31 | 10 | 0 | | |
| RR167_085_Brain_S90 | SAMN45838137 | SRR31744727 | N2302789 | 5 | Control | Brain | 4784130 | 1744780 | 0 | 13 | NA | 0 | | |
| RR167_086_Lung_S1 | SAMN45838126 | SRR31744726 | N2305105 | 6 | Control | Lung | 14295714 | 2410126 | 0 | 0 | 0 | 0 | 0 | |
| RR167_087_Liver_S2 | SAMN45838127 | SRR31744724 | N2305105 | 6 | Control | Liver | 5749486 | 967930 | 1 | 6 | 6 | 0 | | |
| RR167_088_Spleen_S3 | SAMN45838128 | SRR31744723 | N2305105 | 6 | Control | Spleen | 1824424 | 472574 | 0 | 39 | NA | 0 | | |
| RR167_089_Brain_S4 | SAMN45838129 | SRR31744722 | N2305105 | 6 | Control | Brain | 4692484 | 1973504 | 1 | 24 | 24 | 0 | | |
| Water_1_S91 | SAMN45838138 | SRR31744721 | NA | NA | Control | Water Control | 164022 | 1038 | 0 | 0 | 0 | 0 | | |

| Viral Genus | Species | Segment | Host | Genbank Accession Number |
|---|---|---|---|---|
| Bandavirus | Bhanja virus | L | Tick | NC_027140 |
| | | M | | NC_027141 |
| | | S | | NC_027142 |
| | Guertu virus | L | Tick | NC_043611 |
| | | M | | NC_043609 |
| | | S | | NC_043610 |
| | Heartland virus | L | Human | NC_024495 |
| | | M | | NC_024494 |
| | | S | | NC_024496 |
| | Hunter Island virus | L | Tick | NC_027717 |
| | | M | | NC_027715 |
| | | S | | NC_027716 |
| | Lone star virus | L | Tick | NC_021242 |
| | | M | | NC_021243 |
| | | S | | NC_021244 |
| | Razdan virus | L | Tick | NC_022630 |
| | | M | | NC_022631 |
| | | S | | NC_022632 |
| | SFTS virus | L | Human | NC_043450 |
| | | | | NC_018136 |
| | | M | | NC_043451 |
| | | | | NC_018138 |
| | | S | | NC_043452 |
| | | | | NC_018137 |
| Beidivirus | Dipteran beidivirus | L | Housefly | NC_032158 |
| | | M | | NC_032159 |
| Goukovirus | Cumuto goukovirus | L | Mosquito | NC_043045 |
| | | M | | NC_043046 |
| Ixovirus | Scapularis ixovirus | L | Tick | NC_055432 |
| Phlebovirus | Oriximina phlebovirus | L | Uknown | NC_055303 |
| | | M | | NC_055304 |
| | | S | | NC_055305 |
| | | L | | NC_014397 |

| | | | | |
|---|---|---|---|---|
| | Rift Valley fever phlebovirus | M | Human | NC_014396 |
| | | S | | NC_014395 |
| | Sicilian phlebovirus | L | Human | NC_015412 |
| | | M | | NC_015411 |
| | | S | | NC_015413 |
| Uukuvirus | Rukutama uukuvirus | L | Tick | NC_055368 |
| | | M | | NC_055367 |
| | | S | | NC_055366 |
| | Uukuniemi uukuvirus | L | Unknown | NC_005214 |
| | | M | | NC_005220 |
| | | S | | NC_005221 |
| | Zaliv terpeniya uukuvirus | L | Tick | NC_055356 |
| | | M | | NC_055354 |
| | | S | | NC_055355 |

| Animal ID | PCR Ct |
|---|---|
| NJ-1 | 21.785 |
| NJ-2 | NA |
| NJ-3 | NA |
| NJ-4 | NA |
| NJ-5 | NA |
| NJ-6 | NA |
| PA-1 (Bird-1) | 32.455 |
| PA-2 (Bird-2) | NA |
| PA-3 (Bird-3) | 21.881 |
| PA-4 | 32.964 |
| PA-5 | NA |
| PA-6 | NA |
| PA-7 | NA |
| PA-8 | 26.695 |
| PA-9 | 35.316 |
| PA-10 | 24.253 |
| PA-11 | 23.191 |
| PA-12 | 21.37 |
| PA-13 | 24.994 |
| PA-14 | 33.445 |
| PA-15 | 17.882 |
| PA-16 | 34.52 |
| PA-17 | 35.524 |
| PA-18 | 25.585 |
| PA-19 | 20.593 |
| PA-20 | 33.448 |
| PA-21 | 21.37 |
| PA-22 | NA |
